# Phosphorylation and dephosphorylation of Ser852 and Ser889 control clustering, localization, and function of PAR-3

**DOI:** 10.1101/2020.05.04.075952

**Authors:** Kazunari Yamashita, Keiko Mizuno, Kana Furukawa, Hiroko Hirose, Natsuki Sakurai, Maki Masuda-Hirata, Yoshiko Amano, Tomonori Hirose, Atsushi Suzuki, Shigeo Ohno

## Abstract

Cell polarity is essential for various asymmetric cellular events, where the partitioning defective (PAR) protein, PAR3, plays a unique role as a cellular landmark to establish polarity. In epithelial cells, PAR3 localizes at the subapical border such as the tight junction in vertebrates and functions as an apical determinant. Although there is much information about the regulators of PAR3 localization, the mechanism involved in PAR3 concentration and localization to the specific membrane domain remains an important question to be clarified. In this study, we demonstrate that ASPP2, a stimulator of PAR3 localization, can link PAR3 and protein phosphatase 1 (PP1). The ASPP2–PP1 complex dephosphorylates a novel phosphorylation site, Ser852, of PAR3. Furthermore, Ser852- or Ser889-unphosphorylatable PAR3 mutants form protein clusters and ectopically localize to the lateral membrane. Concomitance of clustering and ectopic localization suggests that PAR3 localization is a consequence of local clustering. We also demonstrate that unphosphorylatable forms of PAR3 are static in molecular turnover and fail to coordinate rapid reconstruction of the tight junction, supporting that both phosphorylated and dephosphorylated states are essential for the functional integrity of PAR3.

**Summary statement:** We show that phosphorylation and dephosphorylation regulate clustering of PAR-3, a cell polarity-regulating factor, and how the clustering regulation affects localization of PAR-3 and cell-cell junction formation.

## Introduction

Cell polarity, one of the basic properties of cells, results in the asymmetric distribution of cell components and is governed by sets of evolutionarily conserved polarity-regulating factors such as the PAR–aPKC system. PAR proteins were first identified in the asymmetric cell division of *C. elegans* zygotes (Kemphues et al., 1988). Mutants of these factors failed in the asymmetric cell division, and some of these proteins were found to asymmetrically localize to the anterior side or the posterior side (Etemad-Moghadam et al., 1995; Tabuse et al., 1998). The PAR–aPKC system functions in various asymmetric biological processes of animals such as asymmetric cell division of stem cells and establishment and maintenance of the asymmetric apical and basolateral membrane domain in epithelial cells (Izumi et al., 1998; Knoblich, 2001; Ohno, 2001; Suzuki and Ohno, 2006; Tepass et al., 2001). Although the downstream molecules of these factors are still under investigation, the basic concept for the mechanism of polarity establishment by the PAR–aPKC system has already been proposed, that is, mutual antagonism and positive feedback enhancement. The apical determinant aPKC phosphorylates the basolateral regulators PAR1 and Lgl and excludes them from the apical domain (Betschinger et al., 2003; Suzuki et al., 2004; Yamanaka et al., 2003). Inversely, PAR1 phosphorylates the apical determinant PAR3 and excludes it from the lateral membrane domain (Benton and St Johnston, 2003). In addition, Lgl inhibits the activity of aPKC (Yamanaka et al., 2003). The essentiality of the positive feedback loop that enables self-recruitment of apical determinants was demonstrated in aPKC and the Crumbs complex, another apical determinant protein complex, using both computer simulations and in vivo experiments (Fletcher et al., 2012).

In epithelial cells, the PAR3–aPKC–PAR6–Cdc42 complex (the PAR complex) functions as an apical determinant. PAR3 localizes at the subapical membrane domain, the tight junction in mammalian epithelial cells, and the adherens junction in *Drosophila*, whereas aPKC and PAR6 localize to both the tight junction and the apical membrane domain (Hirose et al., 2002; Morais-de-Sa et al., 2010). PAR3 localizes to primordial adherens junctions prior to other PAR complex components (Suzuki et al., 2002) and plays a unique role in the PAR complex in determining the initial formation of the PAR complex at the cell–cell contact region that becomes the subapical region as a molecular landmark after polarity establishment. All of these observations, combined with other observations on the formation of the tight junction in epithelial cells, implicate the importance of PAR3 localization at a specific membrane domain during polarity establishment (Chen and Macara, 2005; Horikoshi et al., 2009).

PAR3 does not have the transmembrane domain, but it can interact with transmembrane proteins such as JAM (Ebnet et al., 2001) and E-cadherin (Harris and Peifer, 2005) and can also interact with lipids such as phosphatidic acid and phosphoinositides (Horikoshi et al., 2011; Krahn et al., 2010; Yu and Harris, 2012). These interactions can anchor PAR3 to the plasma membrane. The N-terminal oligomerization domain is essential for PAR3 localization to the tight junction region (Mizuno et al., 2003). The PAR3-binding protein ASPP2 is also required for the localization of PAR3 (Cong et al., 2010). Another study demonstrated that phosphorylations of Ser151 (corresponding to Ser144 of mammals) and Ser1085 (corresponding to Ser889 of mammals) in Bazooka/PAR3 by PAR1 result in the exclusion of Bazooka/PAR3 from the lateral membrane in Drosophila epithelial cells (Benton and St Johnston, 2003; Hurd et al., 2003). However, how these mechanisms account for the regulation of PAR3 localization and how these mechanisms relate to each other remain unclear.

In this study, we demonstrate that the ASPP2–PP1 complex dephosphorylates the novel phosphorylation site Ser852 of PAR3 and that dephosphorylation of Ser852 or Ser889 promotes cluster formation and accumulation of PAR3. This phosphorylation–dephosphorylation cycle could ensure the accumulation and turnover of PAR3, which is important for the rapid recruitment of PAR3 to the specific membrane domains where PAR3 acts as a landmark for other polarity regulators.

## Results

### The interaction between PP1 and PAR3 is mediated by ASPP2 and is essential for proper PAR3 localization

As described above, ASPP2 is required for the localization of PAR3 to the tight junction (Cong et al., 2010). In addition to PAR3, ASPP2 can interact with a variety of proteins, including p53. This protein had also been identified as a subunit of protein phosphatase 1α (PP1α), suggesting that PAR3 can form the protein complex with ASPP2 and PP1 (Helps et al., 1995). Consistently, another study demonstrated that PAR3 interacts with PP1α (Traweger et al., 2008). We hypothesized that ASPP2 would bridge the interaction between PAR3 and PP1α and confirmed that PP1α was coimmunoprecipitated with PAR3 (Fig. 1A). By knocking down ASPP2, coimmunoprecipitated PP1α was significantly decreased (Fig. 1A). To investigate the significance of the ASPP2–PP1α association on the regulation of PAR3, we adopted ASPP2 mutants lacking the interaction with PP1α. ASPP2 harbors the evolutionarily conserved RVKF stretch, which matches the PP1-binding consensus sequence, in the PP1-binding domain (Fig. 1B; Fig. S1A) (Egloff et al., 1997). We generated several mutants, including the previously reported ASPP2-REVD mutant (R921E and V922D) (Liu et al., 2011). This mutant was severely defected in the interaction with PP1α (Fig. 1C). YAP and LATS, Hippo pathway factors, have been reported to interact with ASPP2. YAP binds to the proline-rich domain and the SH3 domain, and LATS binds to the PP1α-binding domain of ASPP2 (Rotem et al., 2007; Vigneron et al., 2010). Although the ASPP2 mutant lacking the C-terminus (ASPP2-del3) was impaired to interact with YAP and LATS2, the REVD mutation did not affect the interaction with YAP and LATS2 (Fig. 1C,D). This supports that the REVD mutation does not disrupt functions of ASPP2 other than binding to PP1α. We next expressed intact ASPP2 and ASPP2-REVD mutant to previously established ASPP2-knockdown MDCK cells (Cong et al., 2010). We observed that re-expression of ASPP2 rescued the cell–cell localization of PAR3, whereas the expression of the ASPP2-REVD mutant had no significant effect on PAR3 localization (Fig. 1E,F; Fig. S1B). These results indicate that ASPP2 can act as an adaptor linking PP1 to PAR3 and suggest that this function is involved in the proper cell–cell localization of PAR3.

**Fig. 1.**
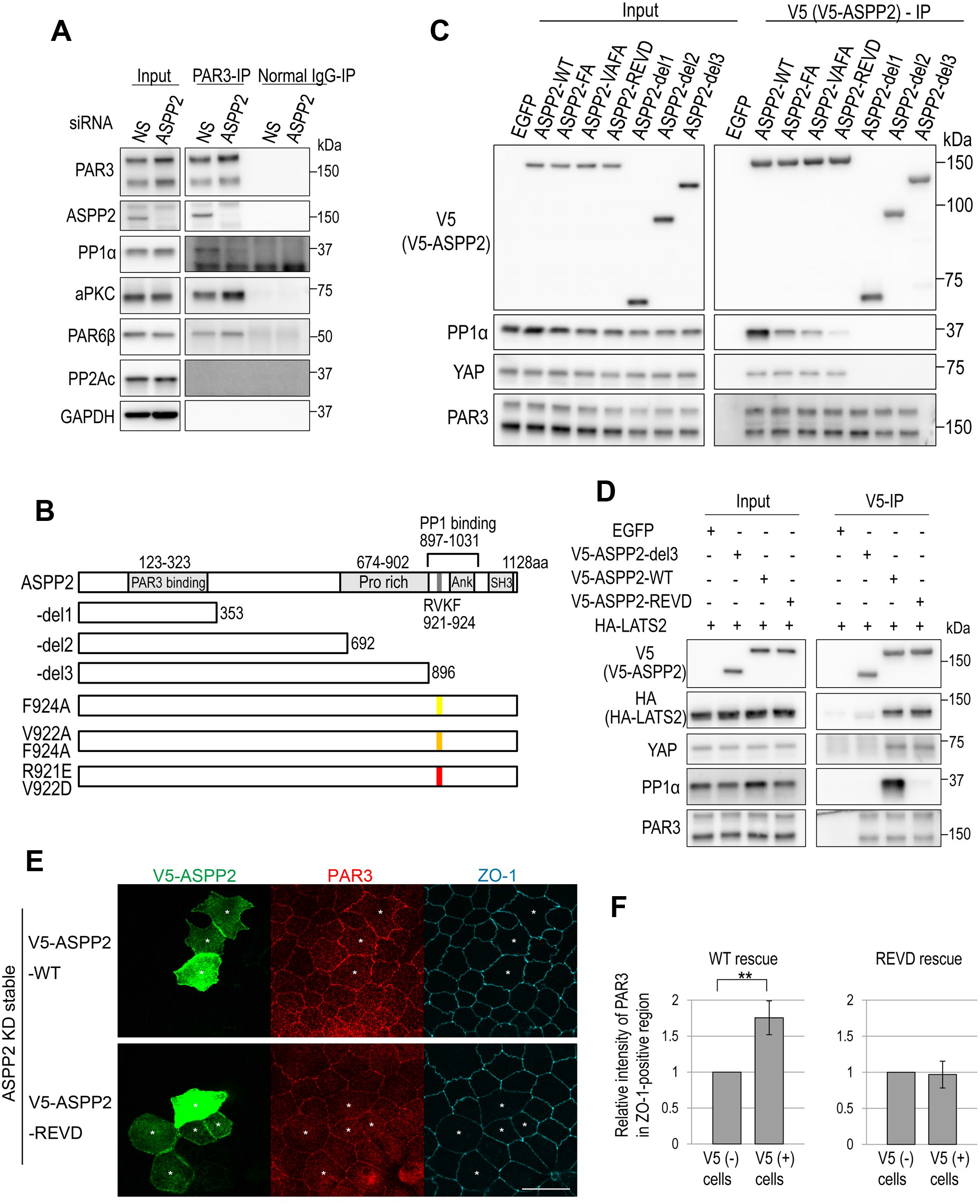
ASPP2 is necessary for the interaction between PAR3 and PP1α, and the interaction between ASPP2 and PP1α is necessary for PAR3 localization. (A) MDCK cells were transfected with nonsilencing siRNA and siRNA for ASPP2. Then, immunoprecipitation was performed using anti-PAR3 or normal rabbit IgG. ASPP2 depletion decreased the coimmunoprecipitation of PP1α with PAR3 (long and short). (B) A schematic representation of the domain structure of ASPP2, its deletion mutants, and point mutants. (C) Interactions between ASPP2 mutants and PP1α or YAP were evaluated by immunoprecipitation. ASPP2 mutants were exogenously expressed in HEK293T cells. (D) Interactions between ASPP2 mutants and LATS2 or PP1α were tested by immunoprecipitation. (E) V5-ASPP2-WT or V5-ASPP2-REVD mutants were expressed in the ASPP2-knockdown MDCK cell line. Rescue of PAR3 localization was assessed. Asterisks indicate V5-ASPP2-expressing cells. (F) Quantification of the fluorescence intensity of PAR3-staining in the tight junction region (approximately 20 cells were measured in each sample, and four photos in two independent experiments were quantified (n = 4)). The precise method is described in Fig. S1B. Scale bar represents 20 μm.

### Ser852 of PAR3 is a novel phosphorylation site that acts as a 14-3-3-binding site

The significance of the ASPP2–protein phosphatase interaction implies that dephosphorylation of PAR3 would promote the junctional localization of PAR3. In fact, several studies had already demonstrated that Ser144 and Ser889 are phosphorylated and serve as binding sites for 14-3-3 and that these phosphorylations regulate PAR3 localization and function (Fig. 2A) (Benton and St Johnston, 2003; Hurd et al., 2003). Initially, we investigated whether the phosphorylations of Ser144 or Ser889 were affected by the depletion of ASPP2. However, there were no significant effects on these phosphorylation levels (Fig. 3C,D). Therefore, we explored unidentified 14-3-3-binding phosphorylation sites in PAR3. First, we performed far-western blotting using several His-T7-Xpress (HTX) tag-fused fragments of PAR3 and bacterially produced GST-14-3-3ζ as a probe. Among them, 1-269 aa fragment exhibited weak affinity to 14-3-3ζ, whereas 710-936 aa fragment showed the strongest affinity (Fig. 2B). This result supports that 14-3-3 binds to phosphorylated Ser144 and Ser889 and that 14-3-3 also recognizes other residues in the 710-936 aa fragment in addition to Ser889. Next, the transcriptional isoforms of PAR3 were subjected to far-western blotting. The mouse short isoform PAR3 that lacks 827-856 aa (sPAR3_ΔPB, RefSeq XP_006531595) was cloned in an earlier study (Hirose et al., 2002), and this isoform exhibited weaker affinity to 14-3-3ζ than to sPAR3, suggesting that important amino acid stretches are located in 827-856 aa (Fig. 2C). On this basis, we produced several Ser point mutants and found that Ser852 would be the novel 14-3-3-binding site of PAR3 (Fig. 2D). Ser852 may be conserved among vertebrates and chordates, whereas S144 and S889 are more widely conserved (Fig. 2E; Fig. S2A,B). We raised antibodies recognizing the respective phosphorylations of Ser852 and Ser889 and confirmed the phosphorylation of Ser852 (Fig. S2C). To evaluate the phosphorylation level of Ser144, we adopted the commercially available monoclonal antibody against the 14-3-3-binding consensus motif (K/RXXpSXP) because the sequence around Ser144 is the only stretch matching this consensus in PAR3 of mouse, rat, and dog. In fact, we confirmed that this monoclonal antibody relatively specifically recognized the phosphorylation of Ser144 (Fig. S2D,E). As it has been reported that Ser151 and Ser1085 of *Drosophila* Bazooka/PAR3 are phosphorylated by polarity-regulating kinase PAR-1, we checked whether PAR-1b/MARK2, one of the four mammalian orthologs of PAR-1, phosphorylates Ser852 of PAR3. The amino acid sequence around Ser852 rather matches the consensus sequence of the PAR-1 substrate (Fig. S2F) (Nesic et al., 2010). In an in vitro kinase assay, it was observed that PAR-1b phosphorylated the PAR3 842-876 aa fragment to the extent comparable to that of tau, an established substrate of PAR-1b (Drewes et al., 1997), but it did not phosphorylate the S852A mutant fragment (Fig. 2F). Consistently, overexpression of PAR-1b significantly upregulated the phosphorylation of both Ser144 and Ser852 (Fig. 2G; Fig. S2G). We next evaluated whether endogenous PAR-1 family kinases contribute to the phosphorylation of Ser852. Because siRNA appears to be unsuitable for inhibiting all of the four PAR-1 homologs, we adopted the GFP-tagged MKI-peptide derived from *Helicobacter pylori* CagA, which would inhibit PAR-1 family kinases (Nesic et al., 2010; Saadat et al., 2007). We observed that MKI successfully inhibited the PAR-1b overexpression-mediated phosphorylation of Ser144 and Ser852. However, it did not significantly inhibit the endogenous phosphorylation of Ser852 and other sites (Fig. 2H). Altogether, PAR-1b can phosphorylate both Ser144 and Ser852 of PAR3. However, not only PAR-1 family kinases but also the other kinases may phosphorylate Ser144 and Ser852.

**Fig. 2.**
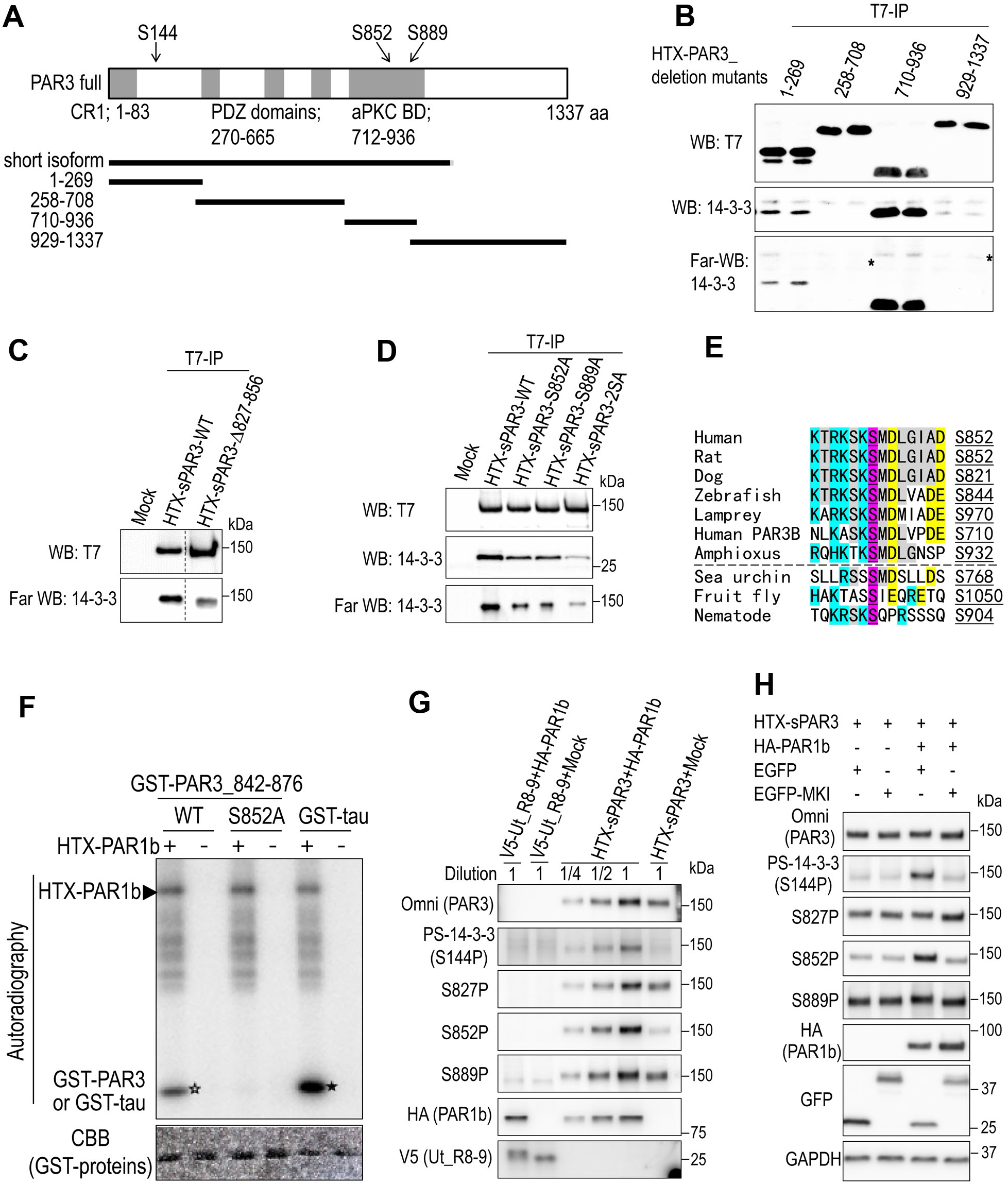
Characterization of PAR3 phosphorylation: Ser852 is a novel phosphorylation site that is important for 14-3-3 binding. (A) A schematic representation of the domain structure of full-length PAR3, sPAR3 (short isoform), and its deletion mutants. (B) Deletion mutants of PAR3 were immunoprecipitated and analyzed by western blotting and far-western blotting using GST-14-3-3ζ as a probe. Asterisks indicate nonspecific signals. Duplicated experiments were performed. (C) PAR3 and PAR3-Δ827-856 deletion mutants were immunoprecipitated and analyzed by far-western blotting. (D) PAR3 and its point mutants were immunoprecipitated and analyzed by western blotting and far-western blotting. Both 852nd and 889th serines were changed to alanines in the 2SA mutant. (E) Sequence alignment around Ser852 of PAR3. Magenta indicates Ser852, and blue, yellow, and gray indicate basic, acidic, and other common amino acids, respectively. Ser852 appears to be conserved among chordates, but not certainly among metazoans. (F) In vitro kinase assay using immunoprecipitated PAR1b as a kinase source and GST-fused PAR3 fragment as a substrate. Open star indicates the phosphorylation of GST-PAR3_842-876, and phosphorylation was not observed in the S852A mutant. GST-tau peptide was used as a positive control (closed star). The arrowhead indicates the autophosphorylation of PAR-1b. (G) HEK293T cells were transfected with each indicated plasmid. Phosphorylations of PAR3 by PAR1b-overexpression were evaluated. Phosphorylation of Ser144 was monitored using the 14-3-3-binding consensus motif antibody. Omni-probe antibody recognizes a part of His-T7-Xpress tags (HTX) as an epitope. The lysate of PAR3- and PAR1b-overexpressed cells (lane 5) was diluted for the quantitative comparison (lanes 3 and 4). Mobility shift of V5-tagged spectrin repeats 8 and 9 of utrophin was used as a positive control of PAR1b-mediated phosphorylation (Yamashita et al., 2010). (H) HEK293T cells were transfected with each indicated plasmid. Promotion of PAR3 phosphorylation by PAR1b-overexpression and the inhibitory effect of MKI on PAR-1b were evaluated. EGFP-MKI almost completely inhibited PAR1b overexpression-mediated phosphorylations (lanes 3 and 4), whereas it did not inhibit the endogenous phosphorylation of PAR3 (lanes 1 and 2).

**Fig. 3.**
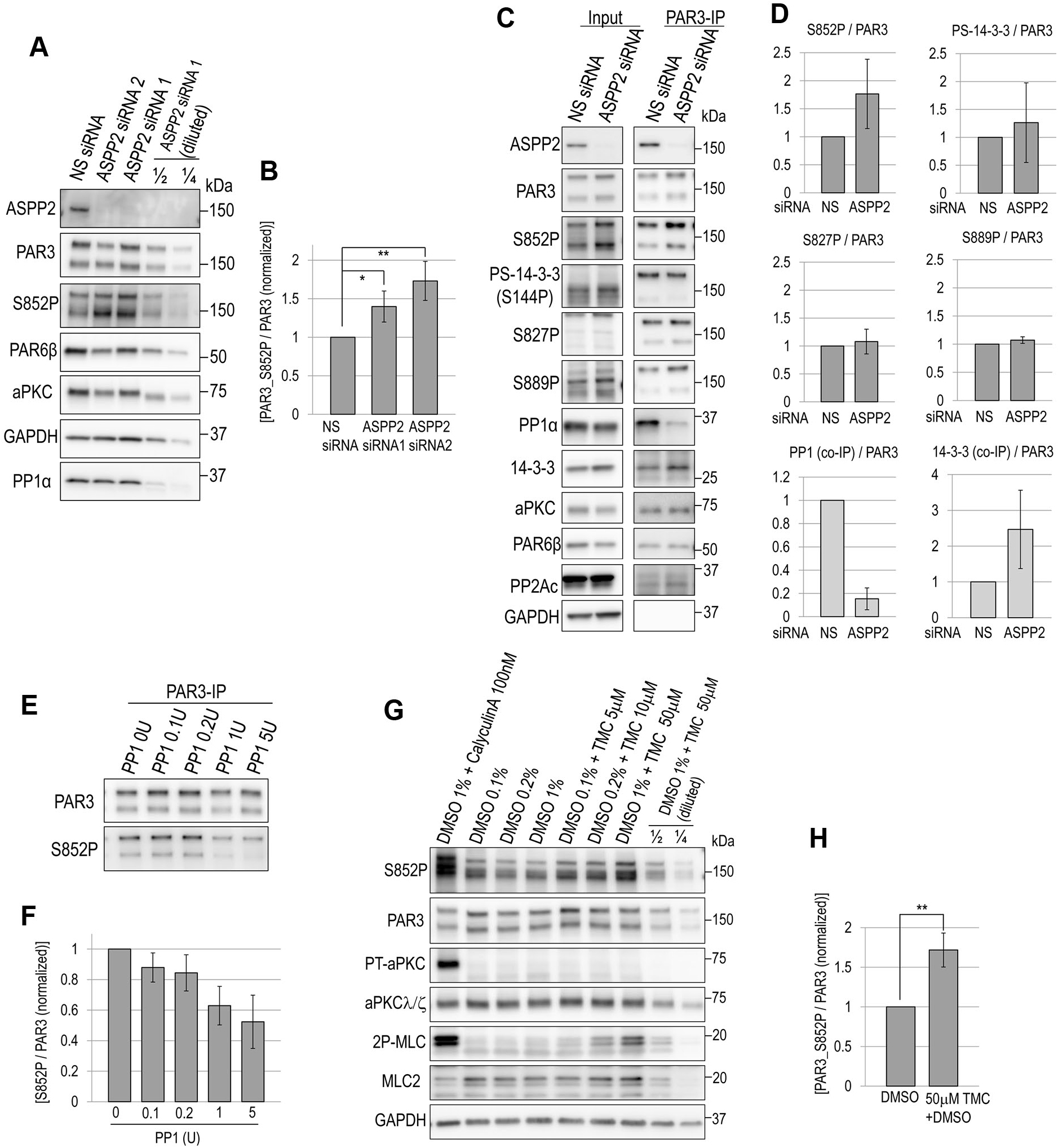
ASPP2 and PPlα negatively regulate the phosphorylation level of Ser852 in PAR3. (A) MDCK cells were transfected with each indicated siRNA, and whole cell lysates were analyzed. The dilution series of the lysate from ASPP2 siRNA1-transfected cells was prepared for the quantification of Ser852 phosphorylation. (B) Quantification of the phosphorylation levels of Ser852 in experiment A (n = 3). The signal intensity of S852P was normalized by the signal intensity of total PAR3. (C) PAR3 was immunoprecipitated from cells that were transfected with each siRNA. Then, the phosphorylation levels and the coimmunoprecipitation with interacting proteins were analyzed. To evaluate the phosphorylation levels of S144, S827, and S889, immunopurified PAR3 was analyzed. (D) Quantification of the phosphorylation levels of PAR3 or coimmunoprecipitation in experiment C (n = 3). (E) PAR3 was immunopurified from MDCK cells, and recombinant PP1α was treated. After incubation, the phosphorylation levels of Ser852 were assayed by western blotting. (F) Quantification of Ser852 phosphorylation in experiment E (n = 3). (G) MDCK cells were treated with calyculin A and tautomycetin (TMC) for 1 h using DMSO as a vehicle. Treatment with both calyculin A and tautomycetin promoted the phosphorylation of PAR3 Ser852 and myosin regulatory light chain (2P-MLC), whereas only calyculin A promoted the phosphorylation of aPKCλ/ζ-Thr412/410. (H) Quantification of Ser852 phosphorylation in experiment G (n = 3).

### The ASPP2–PPlα complex dephosphorylates Ser852 of PAR3

We evaluated whether Ser852 of PAR3 is a dephosphorylation target of the ASPP2–PP1α complex. We observed that depletion of ASPP2 upregulated the phosphorylation level of Ser852 (Fig. 3A,B). We also evaluated the phosphorylation levels of other sites, including Ser827, the target of aPKC (Nagai-Tamai et al., 2002). However, we could not detect any significant change in the phosphorylation levels other than that of Ser852. Owing to the poor specificity of the antibodies, PAR3 immunopurification was required for evaluation (Fig. 3C, lane3 and 4; Fig. 3D). These results suggest that the ASPP2–PP1α complex is involved in the dephosphorylation of Ser852. Next, we assessed the involvement of PP1. The phosphorylation level of Ser852 significantly decreased in a PP1α dose-dependent manner in the in vitro dephosphorylation assay (Fig. 3E,F). PP1α also efficiently dephosphorylated other sites, especially Ser144 and Ser827, suggesting that PP1α can dephosphorylate several sites (Fig. S3A,B). Taken together with the ASPP2-knockdown experiment, PP1α may not be the only factor for the dephosphorylation of Ser144 and Ser827. The significance of PP1 on Ser852 was also confirmed using the PP1-specific inhibitor tautomycetin (Mitsuhashi et al., 2001). Treatment with tautomycetin upregulated the phosphorylation of Ser852 of PAR3 and Thr18/Ser19 of the myosin light chain without affecting Thr412/Thr410 of aPKCλ/ζ and total phospho-Thr, whereas treatment with calyculin A, a nonspecific phosphatase inhibitor, upregulated all of the phosphorylations that we had tested, supporting the specificity of tautomycetin (Fig. 3G,H; Fig. S3C). All of these data support the notion that ASPP2-associated PP1 dephosphorylates Ser852 of PAR3.

**Fig. 4.**
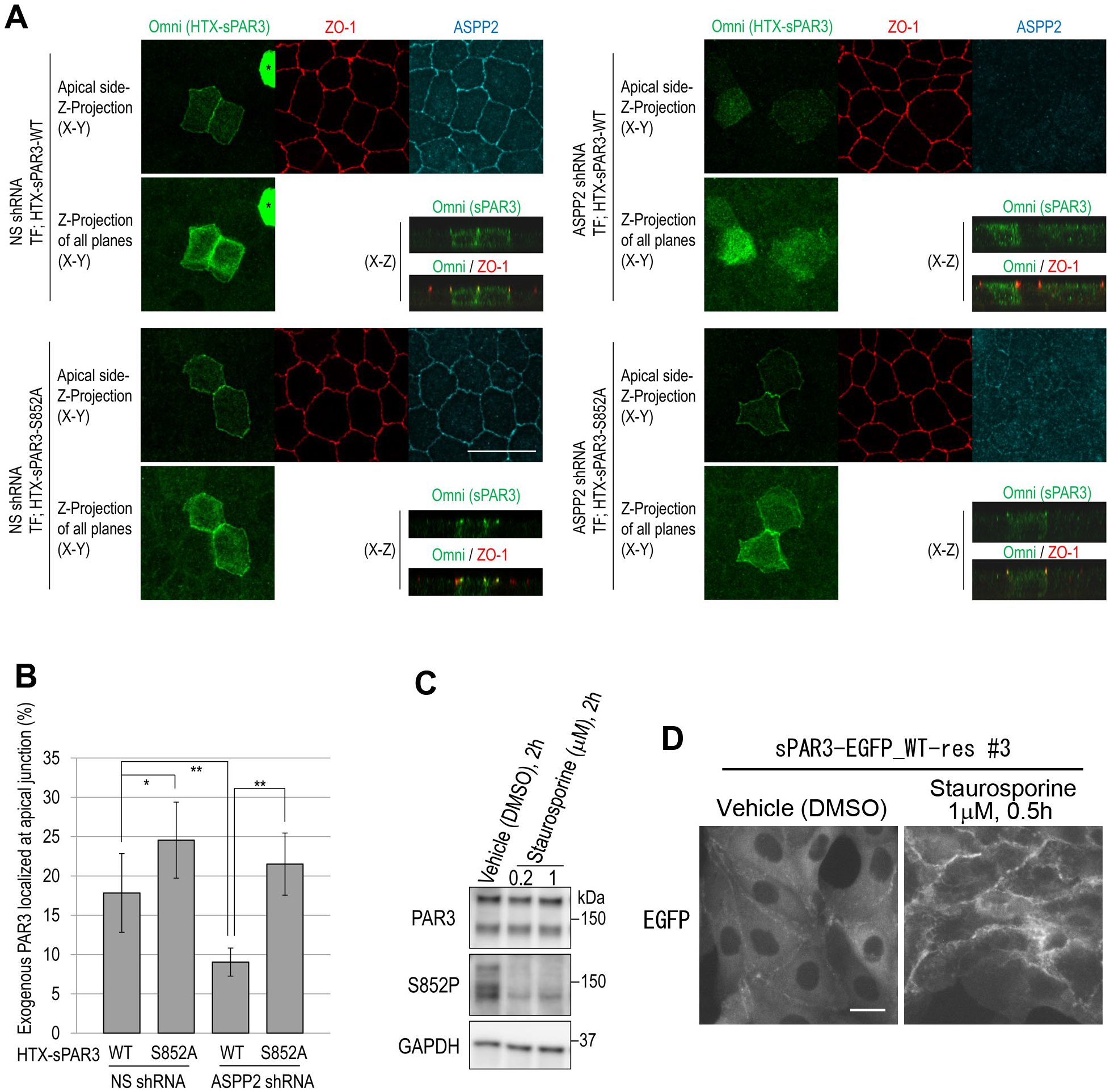
PAR3-S852A strongly localized to the tight junction region irrespective of ASPP2 expression. (A) MDCK cells stably expressing nonsilencing shRNA and MDCK cells stably expressing ASPP2 shRNA were transfected with His-T7-Xpress-tagged sPAR3 wild-type or S852A mutant. Immunofluorescence staining was performed, and samples were observed by confocal microscopy. The percentage of tight junction region-localized PAR3 in total PAR3 was compared. To achieve this, we prepared the sum projection of X–Y sections where ZO-1-staining was positive (left) and the sum projection of all X–Y sections (right). Because most of overexpressed sPAR3 overflowed in the cytoplasm (asterisk), we quantified only the cells expressing low level of sPAR3 by thresholding the intensity. (B) Signal intensities of these images were quantified (approximately 15 cells were measured in each sample, and six photos in two independent experiments were quantified (n = 6)). The precise method is described in Fig. S4. (C) MDCK cells were treated with staurosporine for 2 h at 37°C. Then, the phosphorylation level of Ser852 was evaluated by western blotting. (D) sPAR3-EGFP stably expressing MDCK cells were treated with 1 μM staurosporine for 30 min at 37°C, and localization of sPAR3-EGFP was analyzed. Scale bars represent 20 μm.

### Dephosphorylated PAR3 can localize to the cell–cell junction irrespective of ASPP2 expression

Immunofluorescence experiments showed that both sPAR3-WT and S852A mutant localized to the tight junction region (Fig. 4A,B). Even in ASPP2-knockdown cells, sPAR3-S852A strongly localized to the tight junction region, although the localization of sPAR3-WT was disrupted (Fig. 4A,B). These results indicate the critical role of the ASPP2–PP1α complex in PAR3 localization and dephosphorylation of Ser852. The phosphorylation of Ser852 was strikingly inhibited by the nonspecific kinase inhibitor staurosporine (Fig. 4C), and staurosporine treatment also promoted the cell-cell junction localization of exogenously expressed sPAR3 (Fig. 4D). This result supports the critical importance of the phosphorylation of Ser852 and that dephosphorylation of Ser852 promotes the localization of PAR3 to cell–cell junctions.

### PAR3-S852A and S889A mutants tend to organize ectopic protein clusters and fail to rescue the early step of tight junction formation

Next, we explored the physiological function of Ser852 phosphorylation. For this purpose, we established MDCK cell lines in which endogenous PAR3 was substituted by exogenous EGFP-fused PAR3 and its mutants (PAR3-rescued cells). In addition, S852A, S144A, S889A, S852A/S889A double-mutant, and S144A/S852A/S889A triple-mutant were used in comparing the function of each site. Their expression levels were approximately 10 times greater than the endogenous level (Fig. S5A–C). Using these cell lines, we evaluated junction formation by the calcium switch assay. Through this analysis, we discovered that S852A and S889A mutants were distributed as puncta in the cell–cell junction-disrupted cells (Fig. 5A,D). In several animal species, PAR3 has been observed as puncta, and this structure was believed to be clustered PAR3 (Harris, 2017; Inaba et al., 2015; Tabuse et al., 1998). On this basis, our result suggests that the dephosphorylation of Ser852 or Ser889 tends to cause clusters. Supporting this notion, PAR3 clusters were also observed in PAR3-WT-EGFP-expressing cells treated with staurosporine (Fig. S5D). These clusters merged with aPKC, PAR6β, and ASPP2 but not with GP135, suggesting that these puncta are not vacuolar apical compartments (VACs) and that these clusters contain the PAR–aPKC complex (Fig. S5D,E). Tight junction formation, which was indicated by ZO-1 staining, was not severely disrupted in all PAR3-rescued lines compared to that in the PAR3-knockdown cell line (Fig. 5E; Fig. S5G) (Chen and Macara, 2006; Horikoshi et al., 2009). Although all of these PAR3-rescued lines formed almost complete linear tight junctions at 2 h after the addition of a normal calcium medium (Fig. S5F), the cell lines rescued by S852A, S889A, and S852A/S889A mutants exhibited an obvious defect in tight junction formation at 30 min after the addition of the normal calcium medium compared to that in PAR3-WT-rescued lines (Fig. 5C,E; Fig. S5G). These results indicate that phosphorylations that function to inhibit cluster formation are essential for the early step of tight junction formation. This may be attributed to the prevention of PAR3 recruitment by the clustering-mediated restriction of molecular diffusion because an increased molecular size decreases the diffusion rate (Hofling and Franosch, 2013; Lippincott-Schwartz et al., 2001). Cell lines rescued by PAR3-S144A also exhibited a weaker defect in tight junction formation. This mutation may compromise the tight junction-inducing activity of PAR3 as discussed later.

**Fig. 5.**
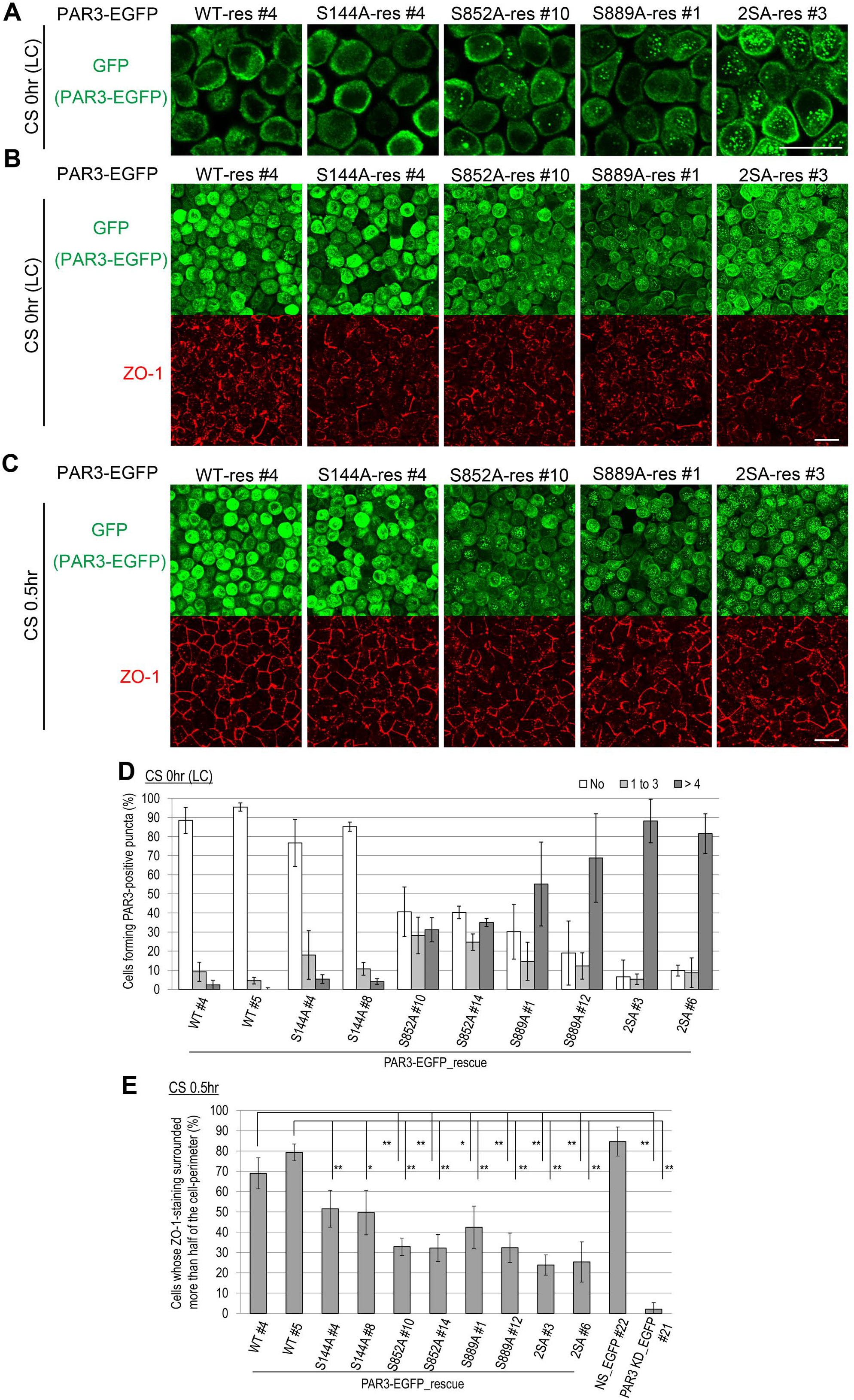
PAR3-S852A and S889A mutants organized ectopic protein clusters and failed to rescue the rapid recovery of ZO-1-staining in calcium switch assay. (A) Wild-type PAR3 and its point mutants were fused with EGFP and stably expressed in the PAR3-knockdown MDCK cell line. These transformant MDCK cell lines were cultured in low-calcium medium for 18 h. Then, immunofluorescence was performed. One confocal section is displayed. (B) Maximum intensity projection (MIP) of confocal sections in (A). (C) After treatment with low-calcium medium, cells were cultured in normal calcium medium for 30 min. The MIP of immunofluorescent confocal sections is displayed. (D) After treatment with low-calcium medium for 18 h, cells positive for punctate PAR3-EGFP staining were counted (at least 50 cells were counted in each sample in one experiment, with three independent experiments (n = 3)). (E) After 30 min of calcium switch, ZO-1-staining was evaluated as an indicator of tight junction maturation. The percentage of cells whose surrounding ZO-1-staining was longer than half the cell perimeter was determined (at least 200 cells were counted in each sample in one experiment, with three independent experiments (n = 3)). More precise quantification is described in Fig. S5G. 2SA indicates S852A/S889A double-mutant. Scale bars represent 20 μm.

### Unphosphorylatable PAR3 mutants exhibited a low turnover rate in a developing cell–cell junction

Cluster formation could alter the kinetics of PAR3. Therefore, we evaluated the turnover rate of PAR3-EGFP at cell–cell junctions by fluorescent recovery after photobleaching (FRAP). We observed that fluorescence was recovered after the bleaching process uniformly throughout the cell–cell contact, suggesting that the newly supplied PAR3-EGFP was primarily derived from the cytoplasm, not from the adjacent plasma membrane (Fig. 6A,B). There were no significant differences in the mobile fraction and the half time of recovery for PAR3-S852A in confluent cultures (see also Discussion). The half time of recovery of PAR3-2SA (852/889) appeared to be prolonged in #6 clone (Fig. 6A,C,E) but was not reproduced in #3 clone (Fig. S6A). However, in developing cell–cell junctions, the half time of recovery of both S852A and 2SA (852/889) mutants was significantly longer than that of wild-type PAR3, although no significant difference was observed in the mobile fraction (Fig. 6B,D,F; Fig. S6B). These results support the notion that the low diffusion rate of clustered PAR3 prevents efficient recruitment of PAR3 to cell–cell junctions.

**Fig. 6.**
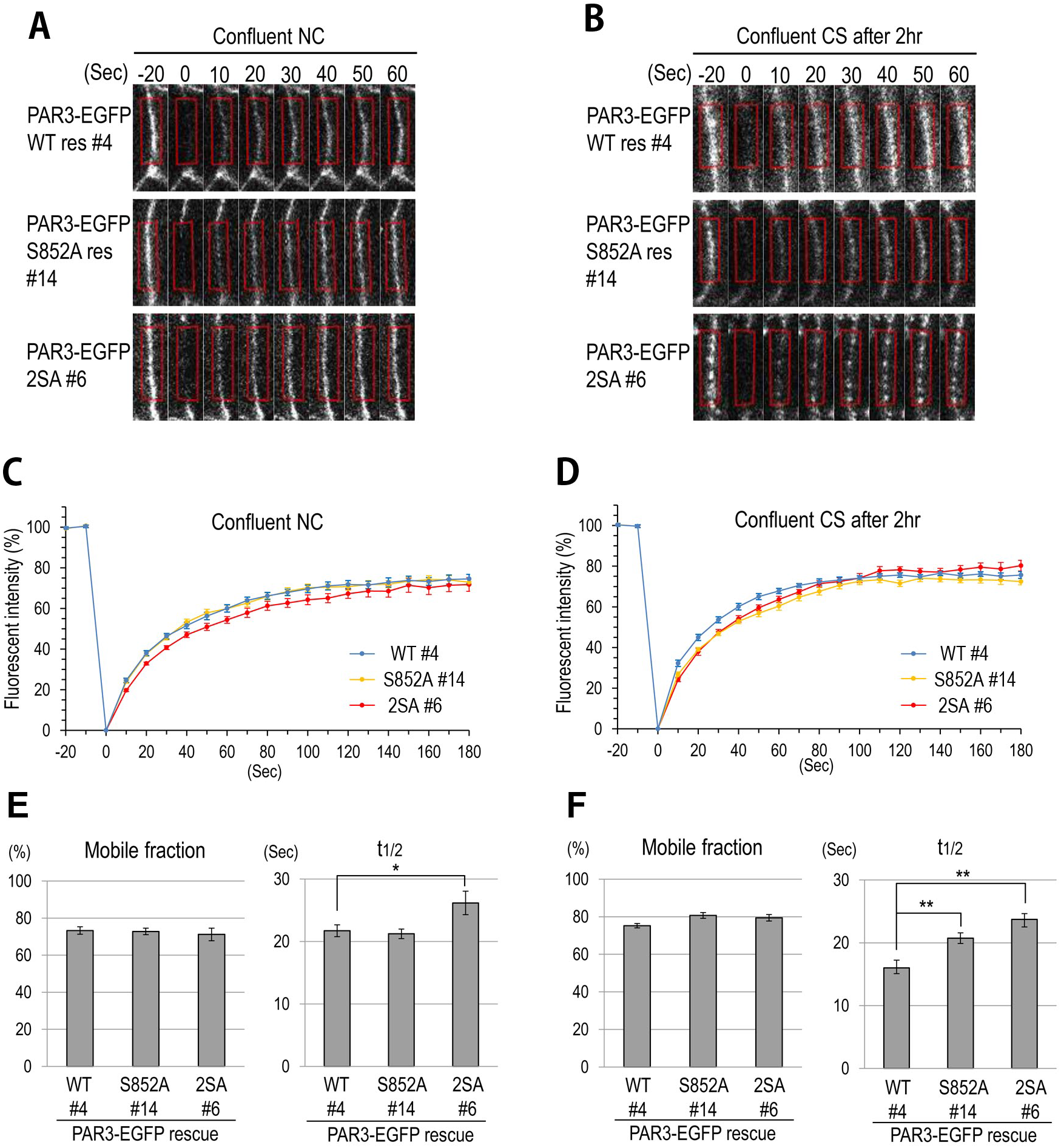
FRAP analysis of PAR3 and its unphosphorylatable mutants in cell–cell junctions. (A) PAR3-EGFP-rescued cell lines were confluently cultured in normal calcium medium and subjected to FRAP assay. Bleaching was performed at time 0. Red rectangles represent ROIs. (B) After calcium switch, cells were cultured for 2 h. Then, they were subjected to FRAP assay. (C) Average values of relative fluorescence intensities in experiment A were plotted (at least 10 ROIs were measured in each sample in three independent experiments). (D) Average values of relative fluorescence intensities in experiment B were plotted. (E) Average values of mobile fraction and half time of recovery in experiment A were plotted. (F) Average values of mobile fraction and half time of recovery in experiment B were plotted.

### S852A, S889A, and S852A/S889A mutants of PAR3 ectopically localize at the lateral membrane and induce the ectopic localization of tight junction components

Ectopic localization of Ser151A and Ser1085A Bazooka/PAR3 mutant proteins to the lateral membrane domain was observed in *Drosophila* (Benton and St Johnston, 2003). In confluent cultures, wild-type PAR3 and its mutants primarily localized around tight junctions. However, S852A, S889A, S852A/S889A, and S144A/S852A/S889A mutants also partially localized to the lateral membrane domain (Fig. 7A). Moreover, in the cells expressing S852A, S889A, and S852A/S889A mutants, tight junction components were ectopically localized to the lateral membrane domain (Fig. 7A,B; Fig. S7A). This suggests that extension of the tight junction region was induced by the ectopically localized PAR3 mutants. When cultured for an additional 2 days, these cells organized the unique intercellular invaginated structures, which contained ZO-1 and appeared like sinuses (Fig. S7B,C). These intercellular sinuses may have been formed by the fracturing of the extended tight junction presumably because of the absence of adherens junctions, which mechanically link cell–cell contacts. This cell morphology is reminiscent of EpCAM-depleted epithelial cells (Salomon et al., 2017). Taken together, these results indicate that dysregulation of PAR3 localization leads to ectopic tight junction formation and morphological defects in the epithelial cell layer.

**Fig. 7.**
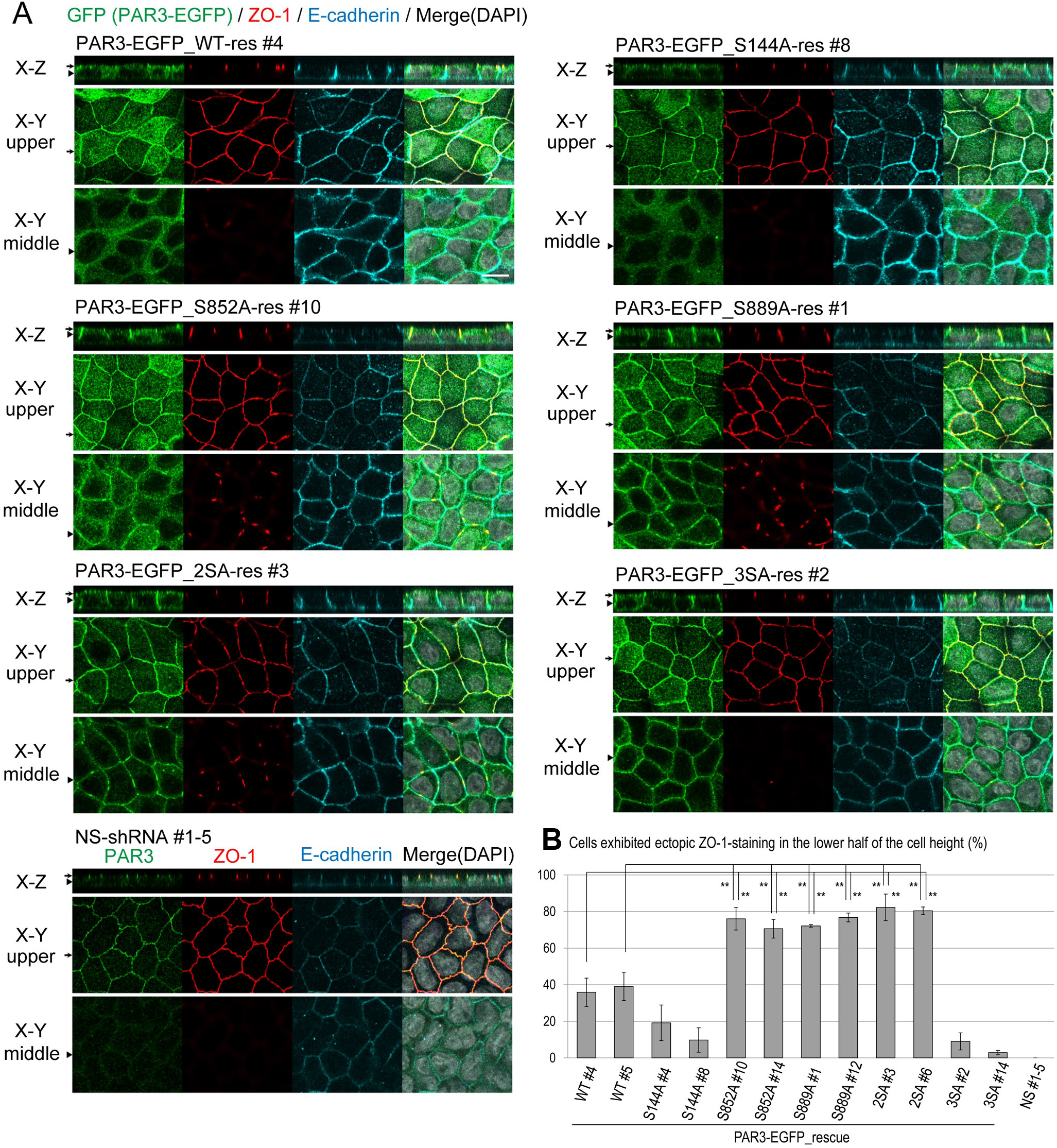
Localization of PAR3 and its unphosphorylatable mutants in PAR3-rescued MDCK cell lines. (A) PAR3-rescued cell lines were cultured for 4 days (reached confluence at day 2). Immunostained samples were analyzed by confocal microscopy. Arrows on reconstituted X–Z sections indicate Z-positions of displayed upper X–Y planes, and arrowheads on the X–Z section indicate Z-positions of displayed middle X–Y planes. Arrows and arrowheads on upper and middle X–Y planes indicate where X–Z sections reconstituted. (B) The percentage of cells exhibiting ectopic ZO-1-staining in the lower half of the cell height was determined (at least 100 cells were counted in each sample in one experiment, with three independent experiments (n = 3)). 2SA and 3SA indicate S852A/S889A double-mutant and S144/S852/S889 triple-mutant, respectively.

## Discussion

In this study, we demonstrated that Ser852 of PAR3, the novel 14-3-3-binding phosphorylation site, was dephosphorylated by the ASPP2-PP1α protein complex and that dephosphorylation of Ser852 and Ser889 promoted the clustering of PAR3. Comparing with the wild-type PAR3, Ser852- and/or Ser889-unphosphorylatable mutants of PAR3 tended to form clusters in the cytoplasm and localize to the plasma membrane when the cell–cell junction was disrupted by calcium depletion (Fig. S5E). Furthermore, these mutants not only localized to the cell–cell contact sites but also mislocalized to the lateral membrane domain under normal culture conditions. Importantly, ectopic clustering and ectopic membrane localization of PAR3 appeared to be concomitant. The results of the previous report from St Johnston laboratory appear to support this notion (Benton and St Johnston, 2003), i.e., unphosphorylatable mutants of Bazooka/PAR3 tended to form clusters compared with the wild-type Bazooka/PAR3, as depicted in Figure 4. Altogether, these data suggest that the so-called “localization” of PAR3 is the consequence of local clustering of PAR3 on the specific plasma membrane domain. The ASPP2–PP1 complex is efficiently recruited to the PAR3 cluster, which harbors several ASPP2-binding sites, and this would further promote ASPP2–PP1-mediated dephosphorylation of PAR3. Our results suggest that this positive feedback loop accumulates PAR3 at the specific membrane domain.

Phosphorylation of Ser852 or Ser889 renders PAR3 diffusive and easily accessible to the newly organized cell–cell junction. When it reaches the cell–cell junction by diffusion, PAR3 may be efficiently dephosphorylated by the ASPP2–PP1 complex, which has been clustered with PAR3, and oligomerize with the cluster. This mechanism may account for the observation that the turnover rates of the wild-type and unphosphorylatable mutant PAR3 were not significantly different on the mature cell–cell junction (Fig. 6E; Fig. S6A). Dephosphorylated PAR3 molecules can concentrate and exert a strong activity that promotes the formation of the tight junction and the apical domain. However, since dephosphorylated form is static in molecular turnover, it fails to coordinate the rapid reconstruction of the tight junction. Therefore, both phosphorylated and dephosphorylated states are essential for the rapid recruitment and the functional integrity of PAR3 (Fig. 8).

**Fig. 8.**
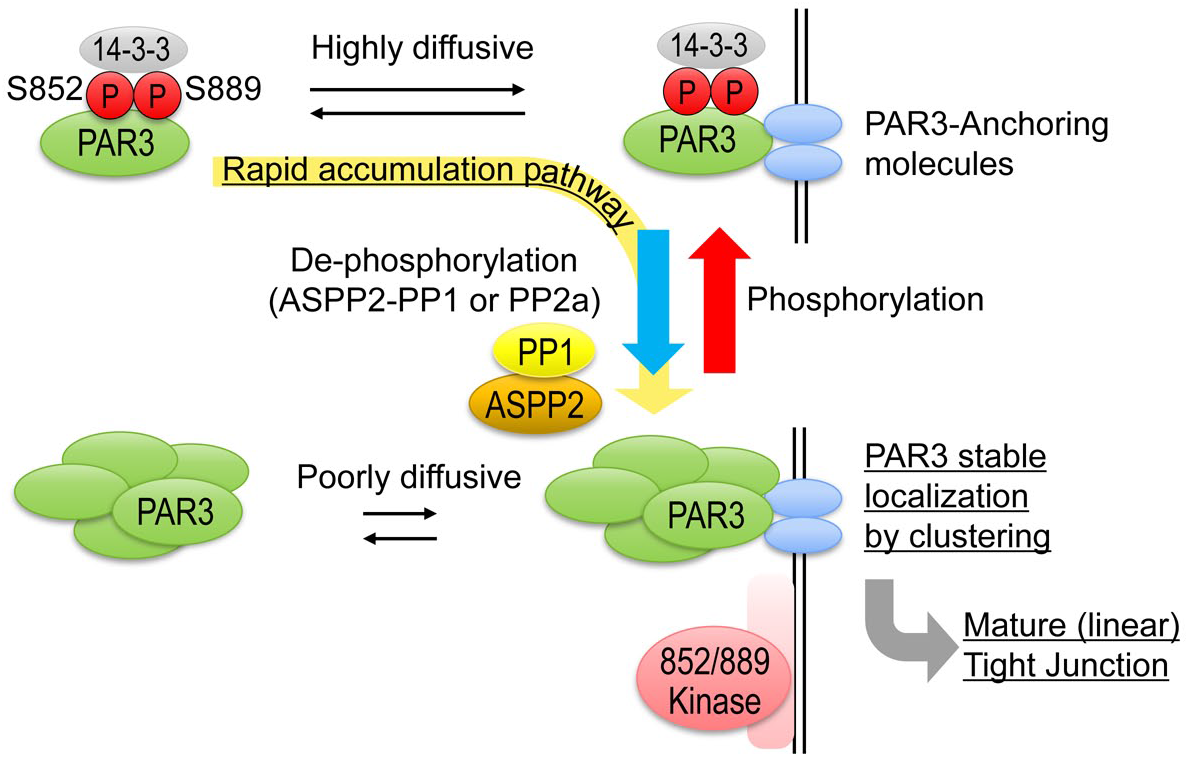
Rapid and proper localization of PAR3 is achieved by the phosphorylation–dephosphorylation cycle. Phosphorylation of Ser852 and Ser889 residues permits the diffusion of PAR3 through association with 14-3-3, whereas their dephosphorylation promotes the clustering of PAR3. Both phosphorylation and dephosphorylation are essential for rapid recruitment and accumulation at specific sites of the membrane such as the cell–cell contact sites (yellow arrow). The balance between kinases localizing at the lateral membrane, possibly PAR1, and phosphatases, including PP1 that is associated with PAR3 through ASPP2, may determine the site where PAR3 accumulates.

On the basis of our observation and previous reports, dephosphorylated PAR3 can localize to several plasma membrane domains (Benton and St Johnston, 2003; Morais-de-Sa et al., 2010). This suggests that the anchoring molecules for PAR3 are not restricted to the tight junction region. Considering this, among the several reported PAR3-binding proteins and lipids, ubiquitously distributed lipids are preferable candidate for the major membrane-anchoring factors. On this basis, Ser852 and Ser889 double-phosphorylated PAR3 can also associate with the membrane-anchoring factors, although the interaction is unstable and transient because of the lack of clustering competency. In our hypothesis, the site where PAR3 clusters is fundamentally important for PAR3 localization. This may be defined by the competition between the phosphorylations by the kinases phosphorylating Ser852 and Ser889 and the dephosphorylation by the ASPP2–PP1 complex engaged to the PAR3–protein cluster. Ser852 kinases may include PAR-1 (Fig. 2F–H). The other kinases, which may also localize to the basolateral membrane domain, should be identified in future studies.

Because the effects of Ser852 and Ser889 phosphorylation on clustering were similar, they appeared to be functionally redundant. However, dephosphorylation of Ser852 was primarily mediated by PP1 in contrast to Ser889, which was reported to be primarily regulated by PP2A (Krahn et al., 2009). Thus, their upstream regulations are different, and the regulations of these sites may be context-dependent. In our observation, the involvement of Ser144 phosphorylation in clustering was not significant. However, the S144A mutant failed to rescue tight junction formation after the calcium switch, and the addition of S144A mutation abrogated the ectopic tight junction-inducing activity of PAR3-S852A/S889A (Fig. 5; Fig. 7). These results suggest that the phosphorylation of Ser144 somehow positively regulates tight junction formation, although the precise mechanisms remain unknown.

The mechanism by which the phosphorylation of Ser852 and Ser889 regulates clustering remains unclear. Ser852 and Ser889 are located in the C-terminal half of PAR3, although the oligomerization domain is located at its N-terminus (Mizuno et al., 2003). The structure of the PAR3 N-terminal domain was revealed by NMR, and the higher order structure of oligomerized PAR3 N-terminal domain was also analyzed. The oligomers showed filamentous structure, which is supposed to be formed by the front-to-back interaction mediated by both type I and type II PB1-like domains of the monomers (Feng et al., 2007; Zhang et al., 2013). Although the PAR3 N-terminal domain can form filamentous oligomers by itself, the front-to-back type of interaction alone cannot appear to organize massive clusters that were observed in cells. Hence, it is plausible that other molecules bind to PAR3 and bridge PAR3 oligomers (Harris, 2017) and that the phosphorylation of Ser852 and Ser889 would be involved in the regulation of this interaction.

Clustering of PAR3 has been broadly observed in several animal species. PAR3 clusters anchored centrosomes to the apical domain in the intestinal cells of *C. elegans* and germline stem cells of male *Drosophila* (Feldman and Priess, 2012; Inaba et al., 2015). In *C. elegans* oocytes, clustering contributes to the efficient transport of PAR3 to the anterior cortex (Rodriguez et al., 2017; Wang et al., 2017). Ser852 is likely conserved among chordates (Fig. 2E). Although the conservation of S852 in other species is unclear, Ser889 appears to be highly conserved among animal species (Fig. S2B). Thus, phosphorylation may be involved in the regulation of PAR3 clustering in several biological processes of various animal species. Interestingly, it has been reported that plasma membrane tension promotes the clustering of PAR3 in *C. elegans* (Wang et al., 2017). In epithelial cells, PAR3 is localized to the cell–cell junction, which is subjected to mechanical stress exerted by circumferential actin belts. On the basis of these facts, it can be speculated that dephosphorylation of PAR3 might be regulated by mechanosensing.

## Materials and methods

### Cell culture, calcium switch, and drug treatment

MDCK II cells and HEK293T cells were cultured in Dulbecco’s modified Eagle medium-low glucose supplemented with 10% fetal bovine serum and 100 U/mL penicillin/streptomycin at 37°C in a humidified atmosphere containing 5% CO_2_. For immunofluorescence of polarized epithelial cells, 1 × 10^5^ MDCK cells were cultured on permeable filters (Transwell 3460, Corning) for 4 days.

Calcium switch assays were performed as previously described (Yamanaka et al., 2006). Briefly, 1 × 10^5^ MDCK cells were cultured for 3 days to reach confluency. These cells were then incubated in a low-calcium (3 μM) medium for 18 h. Then, the medium was changed to a normal growth medium to initiate junction formation. Tautomycetin was obtained from Tocris, and calyculin A was procured from Cell Signaling Technology.

### Expression vectors, small interfering RNAs, transfection, and establishment of transformant cell lines

V5-human ASPP2 full-length and fragments were amplified from V5-ASPP2-SR by PCR (Cong et al., 2010) and were subcloned into pEB6-CAG (Tanaka et al., 1999). sPAR3-SRHis, a His-T7-Xpress-tagged mouse sPAR3-expressing vector, has been described earlier (Mizuno et al., 2003). sPAR3 and long form rat PAR3 were subcloned into pCAG-GS-neo (Izumi et al., 1998; Yamashita et al., 2015) with EGFP sequence to generate sPAR3-EGFP and PAR3-EGFP expression vectors, respectively. The EGFP sequence was amplified by PCR using pEGFP-N1 (Clontech) as a template. All point mutants were generated by PCR-based site-directed mutagenesis. pcDNA-HA-LATS2 was a kind gift from Dr. Hiroshi Sasaki (Ota and Sasaki, 2008). MDCK cells and HEK293T cells were transfected with plasmids using Lipofectamine 2000 and Lipofectamine LTX (Invitrogen), respectively. ASPP2 siRNA1 #2315 and siRNA2 #3326 have been described previously (Cong et al., 2010). ASPP2 siRNA1 was used unless otherwise indicated. MDCK cells were transfected with siRNAs using Lipofectamine RNAiMAX (Invitrogen). The MDCK-transformant clones expressing nonsilencing shRNA or shRNA for ASPP2 have been described previously (Cong et al., 2010). The puromycin-resistant MDCK PAR3 knockdown clone (25a) and the nonsilencing control clone (1-5) were used in establishing PAR3-EGFP-rescued clones and EGFP-expressing control clones (#21 and #22), respectively (Yamanaka et al., 2006). To establish these clones, PAR3-EGFP-pCAG-GS-neo and its point mutants were transfected to PAR3-knockdown cells and selected in the 800 μg/mL G418-containing medium.

### Antibodies

The rabbit anti-ASPP2 antibodies C2AP and C3AP have been described previously (Cong et al., 2010). Anti-phospho-PAR3 Ser827 has also been described previously (Nagai-Tamai et al., 2002). The antibodies specific for PAR3 phosphorylated on Serine 852 and 889 were raised by immunization of rabbits with the keyhole limpet hemocyanin-conjugated phosphopeptides KSKpSMDLGIC and KSSpSLESLQC, respectively, and were affinity-purified. Anti-GST has been described previously (Izumi et al., 1998). Anti-PAR3 (07-330, Upstate), anti-T7 (69522, Novagen), and anti-PP2Ac (05-421, Upstate) were purchased from Merck Millipore. Omni probe, anti-His-T7-Xpress-tag, (sc-7270 and sc-499), anti-PP1α (sc-7482), anti-pan 14-3-3 (sc-629), anti-aPKC (sc-216), anti-PAR6β (sc-67392), anti-ZO-1 (sc-33725), and normal rabbit IgG (sc-2027) were purchased from Santa Cruz Biotechnology. Anti-V5 (R960-25), anti-claudin1 (71-7800), and anti-occludin (71-1500) were procured from Invitrogen. Anti-phospho-aPKCζ Thr410 (9378), anti-myosin light chain 2 (3672), anti-phospho-myosin light chain 2 (3674), anti-phospho-Ser 14-3-3-binding motif (9606), and anti-phospho-threonine (9386) were obtained from Cell Signaling Technology. Anti-aPKC (610176) and anti-E-cadherin (610181) were purchased from BD BioScience. Anti-GAPDH (ab8245) was obtained from abcam, anti-YAP1 (H00010413-M01) was purchased from Abnova, anti-GFP (598) was procured from MBL, and anti-HA (3F10) was obtained from Roche.

### Immunofluorescence and quantification of fluorescent signals

Cells were fixed with 2% paraformaldehyde in PBS and permeabilized with 0.5% Triton X-100 in PBS. After incubation with a primary antibody, cells were stained with Alexa Fluor-conjugated secondary antibodies (Invitrogen). F-actin was stained with rhodamine-phalloidin. Images were obtained using an epifluorescent microscope (AxioImager, Carl Zeiss) or a confocal laser scanning microscope system (LSM700, Carl Zeiss). The ImageJ software was used for the quantification of fluorescent signals. Regions of interest (ROIs) were defined as described in Fig. S1B and Fig. S4.

### Immunoprecipitation and far-western blotting

HEK293T or MDCK cells were lysed in a buffer containing 25 mM Tris-HCl (pH 7.5), 140 mM NaCl, 2.5 mM MgCl_2_, 1 mM ethylene glycol tetraacetic acid, 0.5% Triton X-100, Complete protease inhibitor cocktail (Roche), and PhosSTOP phosphatase inhibitor cocktail (Roche). After centrifugation, the supernatants were subjected to immunoprecipitation with the indicated antibodies, followed by SDS-PAGE and western blotting. His-T7-Xpress-tagged PAR3 mutants were immunoprecipitated by T7 antibody, separated by SDS-PAGE, and then transferred to a polyvinylidene difluoride membrane. For probing by 14-3-3, the membrane was soaked in a denature buffer (50 mM Tris-HCl, pH 8.3, 7 M guanidine, 50 mM dithiothreitol, 2 mM EDTA) for 1 h and then renatured in a renature buffer (20 mM Tris-HCl, pH 7.4, 140 mM NaCl, 4 mM dithiothreitol, 1 mM MgCl_2_, 10 μM ZnCl_2_, 0.1% bovine serum albumin, 0.1% Nonidet P-40) at 4°C for 4 h. After blocking with 4% skim milk in the renature buffer for 4 h, the membrane was incubated with 10 μg/mL of GST-14-3-3ζ in the renature buffer at 4°C overnight and then subjected to immunoblotting using an anti-GST antibody.

### FRAP analysis

Time-lapse imaging was performed using a confocal microscopy system (Axio imager and LSM700, Carl Zeiss) equipped with a 40× dipping objective (Carl Zeiss) and a culture chamber (INUG2-UK, Tokai Hit).

Cells were maintained at 37°C in FluoroBrite DMEM (Gibco) supplemented with 10% FBS under 5% CO_2_ conditions. ROIs were set on cell–cell contacts, where PAR3-EGFP is concentrated. Fluorescence of EGFP was bleached by a 100% power laser and measured by a 0.5% power laser. Images were obtained every 10 s. The fluorescence intensity immediately after bleaching was considered as the background level, and the half time of recovery (t_1/2_) was calculated by curve-fitting to the equation I(t) = I_max_ • (1−e^−kt^), where k = ln2/t_1/2_, using the Solver tool of Excel (Microsoft).

### In vitro kinase assay and in vitro dephosphorylation assay

The kinase assay was performed as previously described (Yamashita et al., 2010). His-T7-Xpress-tagged PAR-1b was overexpressed in COS1 cells and immunoprecipitated by anti-T7 and then used as the kinase source in this experiment. For the in vitro dephosphorylation assay, the substrate PAR3 was immunoprecipitated from confluently cultured MDCK cells using anti-PAR3 antibody. Beads-immobilized substrate was washed with PMP buffer (50 mM HEPES, pH 7.5, 100 mM NaCl, 2 mM DTT, 0.01% Brij 35, 1 mM MnCl_2_), and PP1α (New England Biolab) was added and incubated for 30 min at 30°C. Then, the phosphorylation levels of PAR3 were analyzed by western blotting.

### Statistical analysis

Differences were considered statistically significant when p < 0.05 as assessed using the Student’s *t*-test. Single asterisk and double asterisks denote p < 0.05 and p < 0.01, respectively. The results are presented as mean ± standard deviation (SD).

## Acknowledgments

We thank Dr. H. Sasaki for providing the materials and the members of Ohno laboratories for their helpful comments.

## Author contributions

Conceptualization: K. Yamashita, K. Mizuno, and S. Ohno. Experiments and data analysis: K. Yamashita, K. Mizuno, K. Furukawa, H. Hirose, M. Masuda-Hirata, N. Sakurai, Y. Amano, and A. Suzuki. Funding acquisition: S. Ohno, K. Yamashita, and K. Mizuno. Original draft: K. Yamashita and K. Mizuno. Writing review and editing: T. Hirose and S. Ohno.

